# Bird-assisted sampling of air temperature reveals three-dimensional cooling effects of urban vegetation

**DOI:** 10.64898/2026.10.22.761016

**Authors:** Giacomo Grosso, Luca Nelli, Claudio Carere, Giacomo Dell’Omo

**Author notes:** Corresponding author: Giacomo Grosso, Department of Ecological and Biological Sciences, University of Tuscia, Viterbo, Italy, Address: Largo dell’Università, 01100 Viterbo VT.

## Abstract

Air temperature in cities is highly variable horizontally and vertically, reflecting the complexity of the urban landscape. Yet, studies of urban temperature still depend largely on ground-based meteorological stations and satellite-derived surface temperature products, both of which provide limited information on near-surface air conditions and vertical structure. We tested a bio-assisted monitoring approach to characterise fine-scale three-dimensional patterns of summer air temperature in the city of Rome, Italy, using homing pigeons (*Columba livia*) equipped with lightweight GPS loggers air-temperature sensors. Flight data collected during summer sampling campaigns between 2023 and 2025 revealed pronounced spatial, vertical, and temporal variability in air temperature over the urban area. Modelling results showed that vegetation had a measurable cooling influence within the lower urban atmosphere with grass cover and wooded patches were both associated with lower air temperatures. Thermal contrasts between land-cover types were the strongest at low altitudes and during morning hours, when air above artificial surfaces was up to approximately 5 °C warmer than above vegetated areas. We also observed a buffering effect associated with green areas: air temperature, particularly near wooded vegetation, remained relatively stable up to approximately 300 m from the patch edge before increasing by as much as 3 °C.

These findings suggest that animal-borne sensing can complement conventional monitoring by capturing air temperature variability across urban landscapes and heights, while also helping to identify when and where urban vegetation most effectively mitigates heat exposure. This approach offers new opportunities to support climate-resilient urban planning and targeted heat mitigation strategies.

## 1. Introduction

Cities are highly heterogeneous environments, composed of artificial surfaces, green areas, waterways, and built structures such as buildings of varying heights and street networks, all of which can generate distinct local microclimates. Urban vegetation plays a key role in mitigating urban heat and reducing thermal stress (Saaroni et al. 2018; Schwaab et al. 2021). Vegetation contributes to local cooling through shading, evapotranspiration, and modification of the surface energy balance (Akbari et al. 2001; Aram et al. 2019; Chen et al. 2014). The effectiveness of this cooling depends strongly on spatial configuration and structural characteristics of vegetation, as well on its interactions with the surrounding urban morphology (Masoudi and Tan 2019), which together determine the capacity of green areas to reduce thermal exposure for urban populations (Venter et al. 2020).

Air temperature in cities is shaped by both spatial and temporal factors, including building density, land-cover composition, and distance from the city centre. Built-up areas tend to accumulate and release heat, whereas vegetated areas can mitigate warming through shading and evaporative cooling, with the magnitude of this effect depending on the size, distribution, and canopy density of green spaces (Bowler et al. 2010; Dràbek and Zalesak 2016; Gill et al. 2007). For this reason, the expansion of green infrastructure is widely recognised as a key strategy for mitigating urban overheating and reducing the impacts of extreme heat events in cities (Leal Filho et al. 2021; Marando et al. 2022).

Traditionally, studies of urban thermal environments have relied on ground-based meteorological stations and satellite-derived land surface temperature (LST) products, such as those derived from MODIS (Qiao et al. 2013). Although these datasets are essential for understanding the horizontal structure of urban thermal patterns (Chakraborty and Lee 2019; Clinton and Gong 2013), they have important limitations: ground-based networks often provide sparse spatial coverage, while LST reflects surface temperature rather than air temperatures (Mutiibwa et al. 2015). As a result, the vertical distribution of air temperature within the urban atmosphere remains poorly understood. Airflow and temperature profiles are influenced by building height and density, surface roughness, anthropogenic heat, and local meteorological conditions (Battista et al. 2021; Manoli et al. 2019), making the three-dimensional characterisation of urban microclimates particularly challenging.

To address these observational gaps, emerging bio-logging technologies offer new opportunities to monitor environmental variables at high spatio-temporal resolution (Bridge et al. 2011; Wilmers et al. 2015). Recent studies have explored the use of miniature sensors attached to animals to collect environmental data across heterogeneous landscapes. For example, bats have been used for bio-assisted sampling (BAS) of nocturnal near-surface air temperature in a Middle Eastern city (Chudnovsky et al. 2023), although measurements were restricted to nighttime and very low altitudes. These findings highlight the potential of animal-borne sensors for urban climate monitoring, while also underlining the need for approaches capable of capturing temperature variability across a broader range of altitudes and times of day.

Homing pigeons (*Columba livia*), owing to their ability to fly across urban areas at a wide range of altitudes, represents a promising biological platform for high-resolution microclimate monitoring (Hoff Brait and Antoniosi 2011). By equipping pigeons with lightweight GPS–temperature sensors, continuous measurements of air temperature can be collected along flight trajectories across heterogeneous urban landscapes.

The present study employs a BAS approach to quantify spatio-temporal patterns of summer air temperature across a medium-size European city. Flight data were integrated with Sentinel-2 land-cover information to compare thermal conditions above different surface types, including artificial surfaces, wooded vegetation, and grass-covered areas. Specifically, we investigated how time of day, land cover, altitude, the proportion of surrounding artificial surfaces, and vegetation characteristics influence air temperature. Our primary aim was to evaluate the potential of animal-borne data loggers as a complementary tool for sampling urban air temperature profiles that are difficult to capture using satellite-derived LST products or fixed meteorological stations alone. We hypothesised that urban vegetation mitigates air temperature not only near the surface but also throughout a measurable portion of the lower urban atmosphere. Understanding the magnitude, vertical profile, and temporal evolution of this cooling effect is essential for developing more effective mitigation strategies in urban environments under changing climatic conditions.

## 2. Materials and methods

The study was conducted in the city of Rome, Italy, during the summers of 2023-2025, when air temperatures are high and the cooling influence of green areas is expected to be particularly relevant, especially in Mediterranean environments (Saaroni et al. 2003; Xoplaki et al. 2003). Rome has a Mediterranean climate, with average summer temperatures ranging between 22 and 25 °C and maximum temperatures commonly above 30 °C (Battista et al. 2021).

### 2.1 Data collection

Twenty-seven adult homing pigeons of both sexes were used in the study. Birds were housed in an experimental loft on the northern side of Rome, approximately 14 km from the city centre. To ensure that individuals were familiar with the study area, pigeons underwent a preliminary training phase consisting of multiple releases at progressively distance from the loft. Release points were located both inside and outside the city, at distances ranging from 21 to 56 km from the loft and in different directions relative to it, in order to diversify flight trajectories and sample multiple sectors of the urban area. Before experimental data collection, birds were released from each experimental release point at least three times. Previous studies have shown that repeated releases from the same location can improve the linearity of homing trajectories (Dell’Ariccia et al. 2008). Pigeons were released either individually or in small groups of two or four individuals as part of a separate experiment unrelated to the aims of the present study.

Throughout the training period, individuals wore plasticine dummies of the same weight and shape as the data loggers to accustom them to carrying the GPS units. The dummies were attached to the birds’ backs using a strip of Velcro (Steiner et al. 2000).

Shortly before each experimental release, the dummy was replaced with a GPS data logger (Axy-Trek TechnoSmart Europe, Italy) equipped with an external temperature sensor. The sensor was positioned so that it did not touch the bird’s body and had a fast response time (≤1 s). The device measured 4 x 2 x 1 cm and weighted 12 g, corresponding to 2.8% of male body mass and 3.4% of female body mass in the pigeons used in the study. The device recorded GPS positions and air temperature at 1 Hz. When pigeons returned to the loft, the device was removed for data download and replaced with the dummy.

Land-cover was derived from a supervised classification of Sentinel-2 satellite imagery at 10 × 10 m spatial resolution (European Space Agency; Drusch et al. 2012), implemented using the Semi-Automatic Classification plug-in in QGIS (Congedo 2021). The image used for the classification referred to the summer of 2024, the middle year of the 2023-2025 sampling period. This image was considered representative of land-cover conditions across the study period, as major changes in urban green areas were not expected over this short time frame. For each GPS point, the corresponding land-cover class was extracted from the classified raster. Land-cover types within the study area (450.28 km²) were grouped into broad classes: artificial surfaces (AR, 53.0 %), wooded vegetation (WO, 8.3 %), grass vegetation (GR, 38.5 %). Water, including the river and other water bodies, was not considered in the analysis because it represented less than 0.2% of the study area.

Additional variables were derived for each GPS point, including the percentage of each land cover within 100 m and 500 m buffers (Knerr et al. 2019), the distance from each land-cover class, and the area, in hectares, of the land cover patch overflown by the bird.

Flight speed, derived from the GPS sensor, was included as a covariate to account for possible airflow-related effects on temperature measurements. Increased airflow around the sensor can enhance convective air exchange and reduce radiative heating, whereas low-ventilation conditions can increase radiative warming of the sensor and lead to an overestimation of air temperature (Nakamura and Mahrt 2005). Wind speed and direction could not be explicitly accounted for because sensor was not equipped with an anemometer. In addition, a Digital Elevation Model (DEM) (Tarquini and Nannipieri 2017) of the study area was incorporated to account for altitudinal effects and terrain-related variability in air temperature

### 2.2 Statistical analysis

For the statistical analyses, we retained only active flight segments occurring within a 12-km radius from the city centre. This circular area was chosen because the densely urbanised core of Rome, which is broadly delimited by a ring road highway, located approximately 12 km from the geometric city centre. This area includes the highest densities of built-up surfaces and major urban parks, making it suitable for assessing variation in air temperature across contrasting urban land-cover types.

Prior to model fitting, the dataset was screened for outliers (Andersen 2008 Bjørneraas et al. 2010). GPS points associated with flight speeds below 10 km h⁻¹ (Gagliardo et al. 2011) were classified as resting or non-active flight points and excluded from the analysis (Fig. S1). This threshold was applied to minimize potential biases in temperature measurements associated with stationary or low-mobility conditions, during which the thermometer could be affected by direct solar radiation and reduced convective cooling, leading to overestimation of ambient air temperature (Teichmann et al. 2024).

Habitat use was evaluated by comparing the GPS locations with land-cover availability within the study area, in order to assess whether pigeons flew over specific land-cover classes more or less frequently than expected based on their availability. Habitat availability was quantified as the proportional area (km²) of each land-cover class within the study area, while land-cover use was quantified as the number of GPS points falling within each class. Expected frequencies were obtained by multiplying the total number of GPS locations by the proportional availability of each land-cover class. A chi-square goodness-of-fit test was used to assess whether land-cover use differed from availability. To quantify the direction and magnitude of selection, selection ratios (SR) were calculated as the proportion of used locations divided by the proportion of available habitat for each land-cover class. Values of SR > 1 indicate positive selection, whereas SR < 1 indicate avoidance. Standardized residuals were computed to identify which habitat categories contributed most to deviations from random use.

To quantify the environmental and flight-related drivers of air temperature recorded during pigeon flights, we fitted generalised additive mixed models (GAMMs). Air temperature recorded by the data loggers was used as the response variable, while candidate explanatory variables included time of day, terrain elevation derived from the Digital Elevation Model (DEM), flight altitude above ground, flight speed, land-cover class, distance to each land-cover class, patch size of the overflown land-cover class, and the percentage cover of each land-cover class within 100 m and 500 m buffers. Date was included as a random effect to account for day-to-day variation in atmospheric conditions.

Models were fitted using the bam() function in the mgcv package in R (Wood 2017). Smoothing parameters were estimated using fast restricted maximum likelihood (fREML), which provides stable estimation of smooth terms and helps reduce overfitting. The final model was selected from a full model containing all candidate variables and biologically relevant interactions, using backward selection guided by Akaike Information Criterion (AIC) values (Bolker et al. 2009; Zuur et al. 2009).

To assess the robustness of model performance to sampling variation, we used a bootstrap resampling procedure at the flight level. Subsamples of increasing size, from 500 to 25,000 observations in increments of 1,000, were drawn with replacement from the full dataset. For each subsample size, the model was fitted 50 times using the same structure as the final GAMM, and the coefficient of determination (R²) was computed for each iteration. The mean and standard deviation of R² across repetitions were then calculated to evaluate the stability of model performance.

To evaluate the predictive capacity of the model, a cross-validation procedure was performed. The data were randomly divided into a training set (80%) and a test set (20%). Predictive performance was evaluated by calculating the root mean square error (RMSE), mean absolute error (MAE) and coefficient of determination R² (Hastie et al. 2009; Willmott and Matsuura 2005).

After fitting the GAMM, temperature differences among land-cover classes were quantified by computing pairwise comparisons between the land-cover–specific smooths of hour of day and altitude. Differences were estimated using the difference_smooths() function from the *gratia* package (Simpson 2024). This approach enabled us to identify the combinations of hour of day and altitude at which predicted air temperature differed among land-cover classes.

### 2.3 Heat mapping scenarios

To map predicted air temperature across the city, on the final GAMM was applied to a regular 100 x100 m spatial grid covering the study area. For each grid cell, all the covariates included in the final model were assigned. Predictions were calculated for each cell using the gg*predict()* function from the *ggeffects* R package (Lüdecke 2018). Scenarios were produced for an average summer day at three representative times of day (9:00, 12:00 and 15:00) and at an elevation of 20 m above ground. Descriptive statistics of predicted air temperature, including minimum, maximum, and mean values, were calculated for the whole study area and separately for each land-cover class (AR, WO, GR).

### 2.4 Identification of priority areas for mitigation measures

By combining the model-predicted air-temperature map with land-cover information, we identified areas of Rome characterised by the high predicted temperatures and limited availability (or lack) of green space, representing potential priority areas for nature-based mitigation measures (Elmarakby and Elkadi 2024).

Thermal hotspots (HT) were identified using a quantile-based threshold: cells with temperature values above the 90^th^ percentile of the overall distribution at 9:00 were classified as high-temperature areas. We also identified cells with low predicted air temperature (LT), defined as those below the 10^th^ percentile.

The local green ratio, with wooded and grass-covered areas considered together, was calculated using a moving window and expressed as the proportion of green areas in the immediate vicinity of each cell. Areas with a green ratio below 10% were considered to have insufficient local green space.

Priority areas for possible intervention were identified by combining thermal hotspots with areas of low green availability (HT_LGREEN). This index allows the spatially explicit identification of urban areas that are both exposed to high predicted temperatures and characterised by limited vegetation cover, supporting the prioritisation of urban greening interventions.

## 3. Results

A total of 27 pigeons were trained and released, yielding 42 complete flight tracks collected during the second half of June across the three study years. Only the portions of the tracks occurring within the study area were retained for the analyses, resulting in a total of 26,280 flight points (Fig. 1, Table S1, Table S3).

**Fig. 1.**
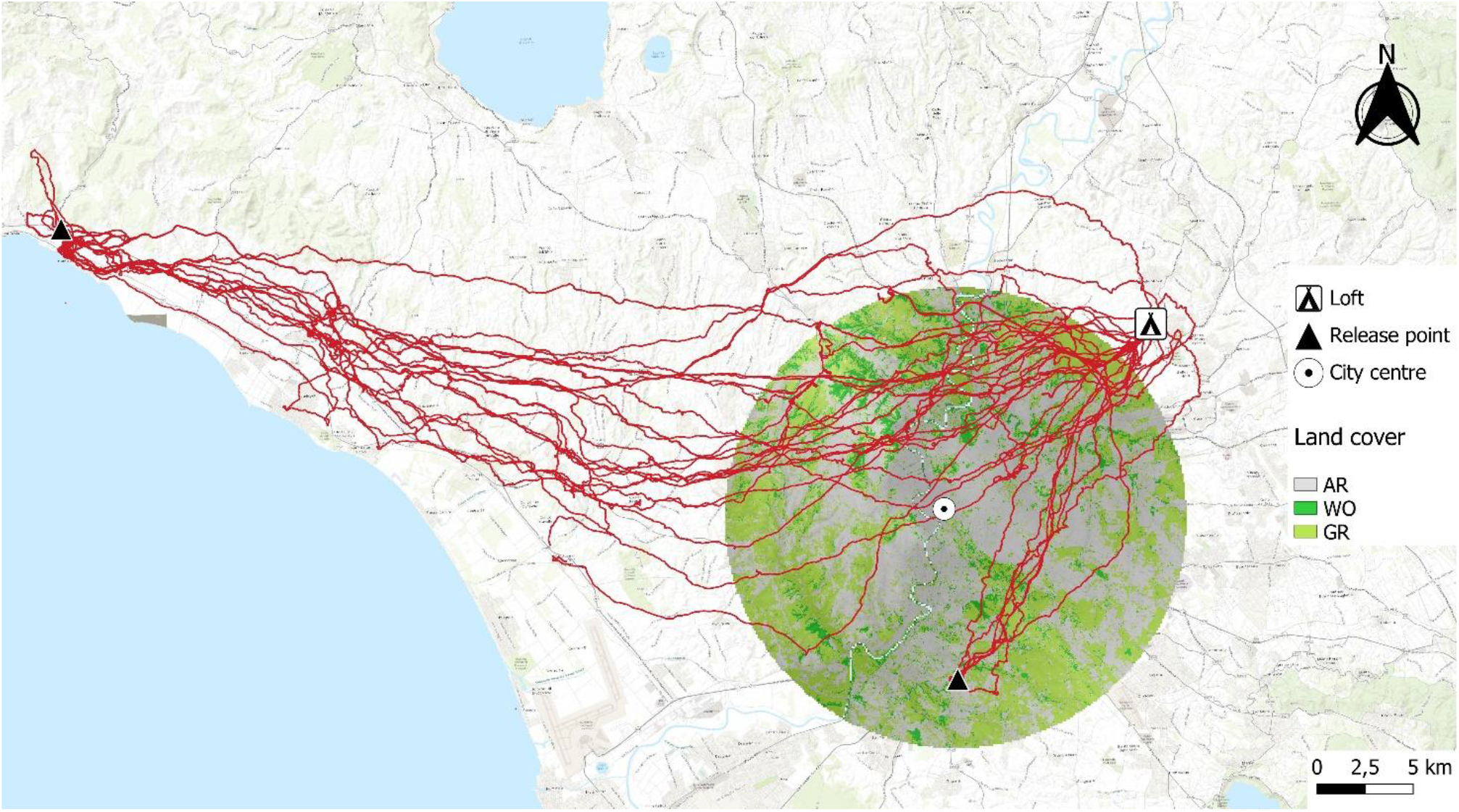
Flight paths of homing pigeons released from two distinct locations, one inside and one outside the city of Rome. Release points, loft location, and city centre are indicated. The circle delineates the study area used for the analyses. Land-cover classes are shown in the background

### 3.1 Statistical analysis

The distribution of GPS points differed among land-cover classes, with 16,995 points over artificial surfaces, 7,740 over grass-covered areas, and 1,545 over Wooded vegetation (Table 1). Habitat use differed significantly from availability (χ² = 1405.4, df = 2, p < 0.001).

**Table 1.** Land-cover composition of the study area and distribution of observed sampling points across land-cover classes. The table reports area (km²), percentage cover, observed and expected point counts based on availability, point density (points km⁻²), selection ratios, and Pearson and standardized residuals from the chi-square test assessing differences between habitat use and availability (χ² = 1405.4, df = 2, p < 0.001). Land-cover classes are defined as artificial surfaces (AR), forest and woodland (WO), and semi-natural non-forest areas (GR)

| Land cover | AR | WO | GR |
| --- | --- | --- | --- |
| Observed points | 16995 | 1545 | 7740 |
| Expected points | 13931 | 2192 | 10115 |
| Study area km<sup>2</sup> | 238.73 | 37.58 | 173.33 |
| Study area cover percentage | 53.01 | 8.34 | 38.49 |
| Percentage of use | 64.67 | 5.87 | 29.45 |
| Selection ratio | 1.22 | 0.70 | 0.76 |
| Pearson residuals | 25.96 | -13.81 | -23.62 |

**Table 1** Land-cover composition of the study area and distribution of observed sampling points across land-cover classes.
| Standardized residuals | 37.87 | -14.43 | -30.11 |
| --- | --- | --- | --- |

AR accounted for 53.01% of the study area but contained 64.67% of the GPS locations, corresponding to a SR of 1.22 and a positive standardized residual (37.87). This indicates that artificial surfaces were over-represented along flight trajectories relative to their availability. In contrast, WO represented 8.34% of the available area but only 5.88% of the locations (SR = 0.70; standardized residual = −14.43), while GR covered 38.49% of the area and contained 29.45% of the locations (SR = 0.77; standardized residual = −30.11). Both vegetated land-cover classes were therefore under-represented along flight trajectories relative to their availability (Fig. 2).

**Fig. 2.**
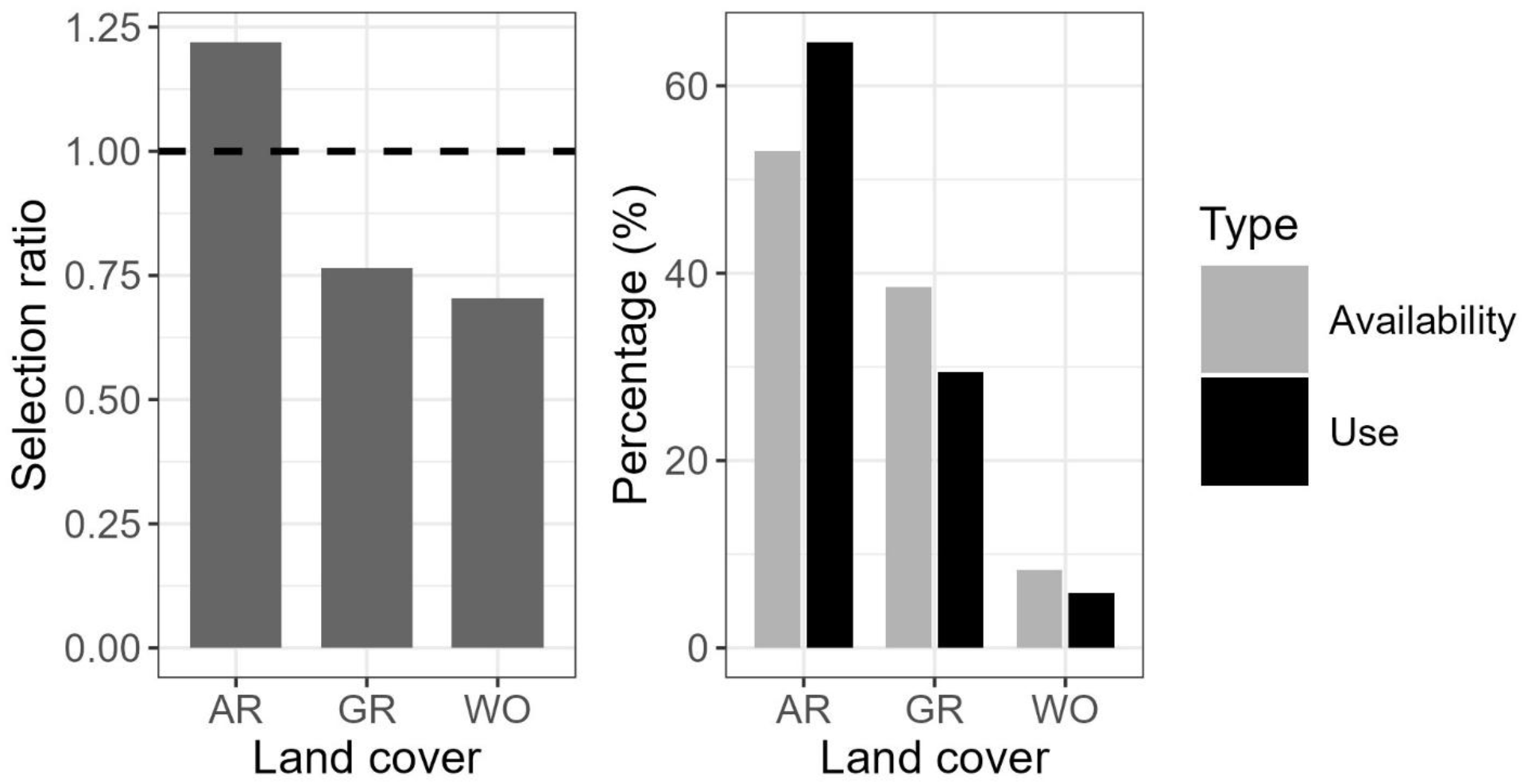
Habitat selection across land-cover classes based on GPS location data. The left panel shows selection ratios for each land-cover class, with the dashed line (selection ratio = 1) indicating proportional use relative to availability. The right panel compares the percentage of habitat availability and observed use. Artificial surfaces (AR) were used disproportionately more than expected based on availability, while forest and woodland (WO) and semi-natural non-forest areas (GR) were used less than expected

The spatial distribution of the tracks showed a lower density of flights in the south-western part of the city. Flight trajectories were shaped by homing behaviour and therefore did not provide a spatially uniform sampling of the urban area. As a consequence, some portions of the city were sampled more intensively than others. However, the objective of this study was not to reconstruct absolute spatial temperature fields, but rather to analyse relative differences in air temperature associated with land cover and terrain characteristics along sampled flight segments. Although behavioural constraints on movement may influence spatial coverage, the modelling framework explicitly accounted for key environmental and flight-related covariates, allowing us to assess temperature variation in relation to land-cover and terrain characteristics within the sampled urban airspace.

During active flights, mean flight speed was 59.5 km h⁻¹ (SD = 13.8), while mean flight altitude was 45.1 m above ground (SD = 23.7). The distributions of flight speed and altitude are shown in Fig. 3. Air temperatures recorded by the sensors ranged from 22.3 to 39.9 °C, with a mean of 30.0 °C (SD = 2.9).

**Fig. 3.**
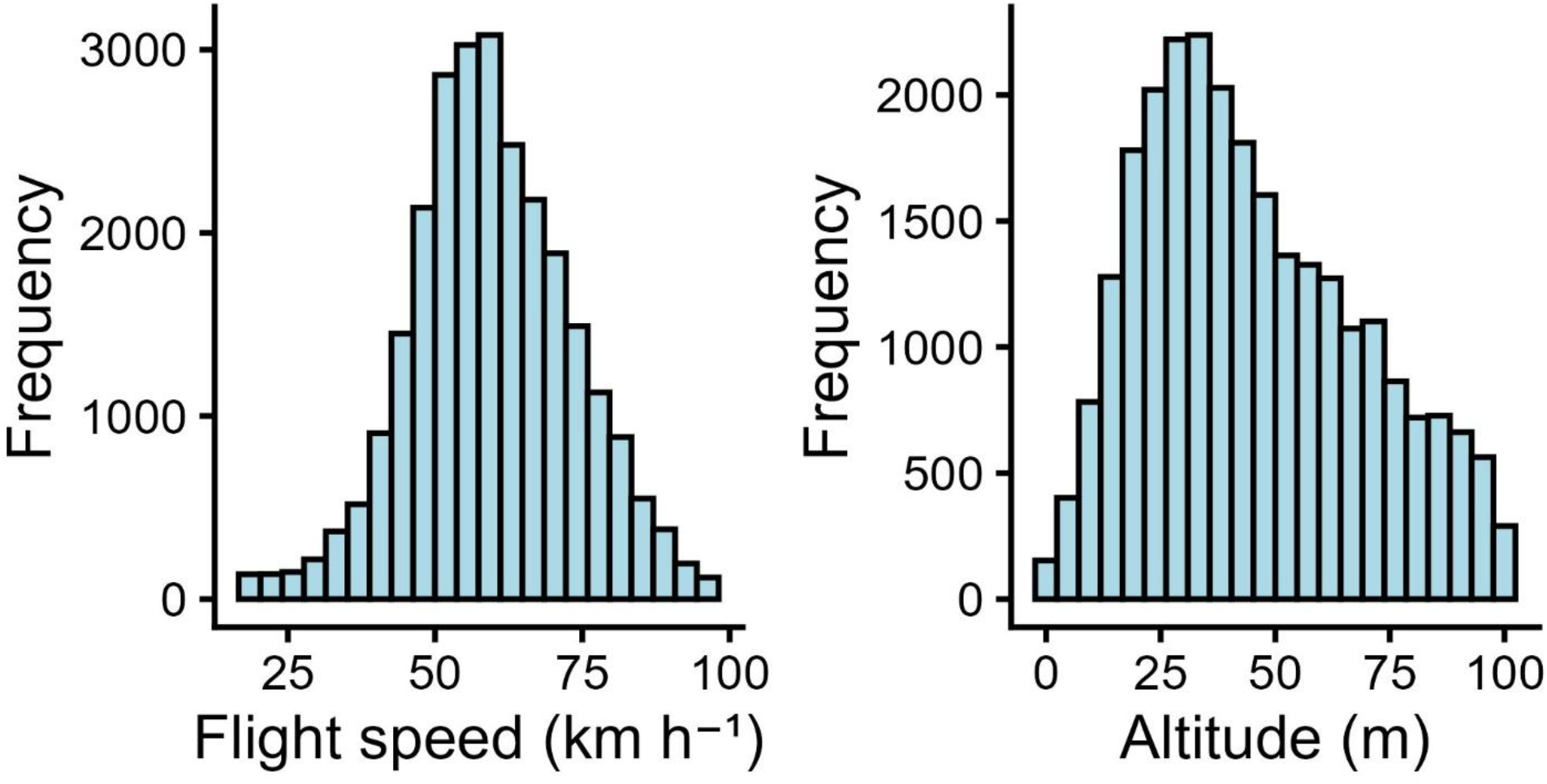
Distribution of flight speed (km h⁻¹) and altitude (m above ground) of pigeons flying over the study area

The GAMM analysis of air temperature recorded during flight as a function of time, land cover and terrain elevation, showed high explanatory power (Table S2, Table S3), accounting for 73.5% of the variance in air temperature (adjusted R² = 0.73, n = 26,280). Model performance assessed through the bootstrap resampling procedure indicated stable explanatory ability while increasing sample sizes (mean R² = 0.73, SD = 0.002). The model also showed good predictive performance in the hold-out validation test (RMSE = 1.48 °C; MAE = 1.04 °C; *R*^2^ = 0.73), indicating that a large proportion of the observed variation in air temperature was explained by the available environmental and urban covariates.

After removal of non-significant terms, namely the percentage of artificial and wooded surfaces, the area of artificial and grass-covered patches, and the distance from artificial surfaces, the final model was selected based on the lowest Akaike Information Criterion (AIC=95,543.4). All terms retained in the final model were statistically significant (p < 0.01).

Parametric terms revealed a significant cooling effect associated with vegetation-related variables. Air temperature decreased with increasing grass cover within a 500 m buffer (β = −0.011 °C per 1% grass cover increase, p < 0.001) (Fig. 4) and with increasing wooded patch area (β = −0.0014 °C per 1 ha in tree-cover increase, p = 0.007) (Fig. S2).

**Fig. 4.**
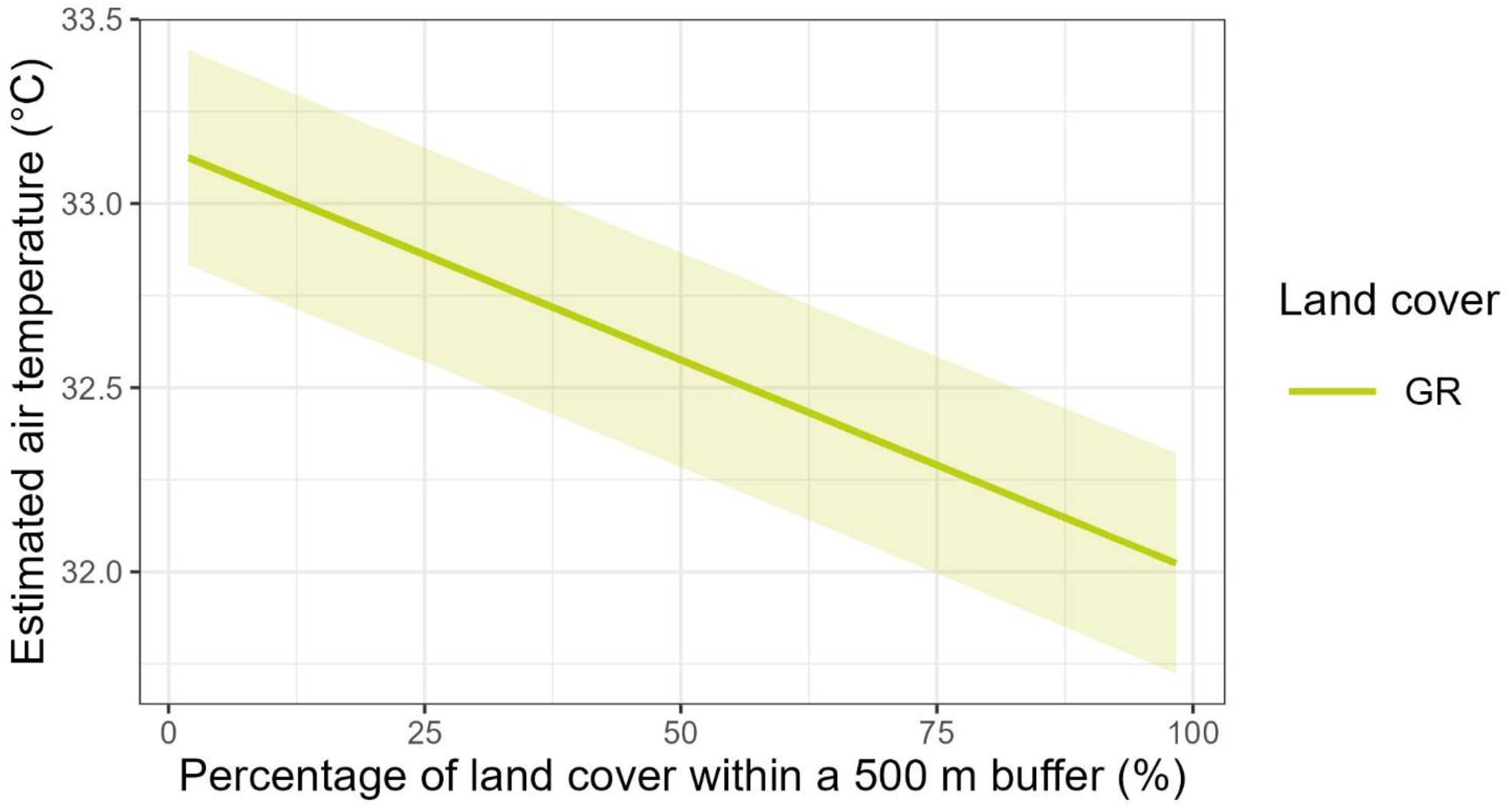
Relationship between predicted air temperature and the percentage of grass-covered areas within a 500 m buffer, as estimated by the final GAMM

Flight speed was retained as a significant covariate (β = −0.0126 °C per 1 km h⁻¹ increase, p < 0.001, Fig. S3), with higher flight speeds associated with slightly lower recorded temperatures. A similar relationship was observed for terrain elevation, with increasing elevation associated with lower air temperature (β = −0.0122 °C per m, p < 0.001) (Fig. S4).

Pairwise comparisons of predicted air temperature among land-cover classes showed marked temporal and vertical differences (Fig. 5). Artificial surfaces were warmer than wooded and grass-covered areas by up to approximately 5 °C, particularly at low altitudes, below 25 m, and during the early part of the day. These differences remained evident up to approximately 75 m. Conversely, during the very early morning and late afternoon, and at higher altitudes, air-temperature differences among land-cover classes tended to decrease, disappearing in some regions of the hour × altitude space and indicating convergence in thermal conditions among land-cover classes. Differences between wooded and grass-covered areas were generally small, although wooded areas tended to show stronger cooling at low altitudes, below 25 m, especially during the early morning.

**Fig. 5.**
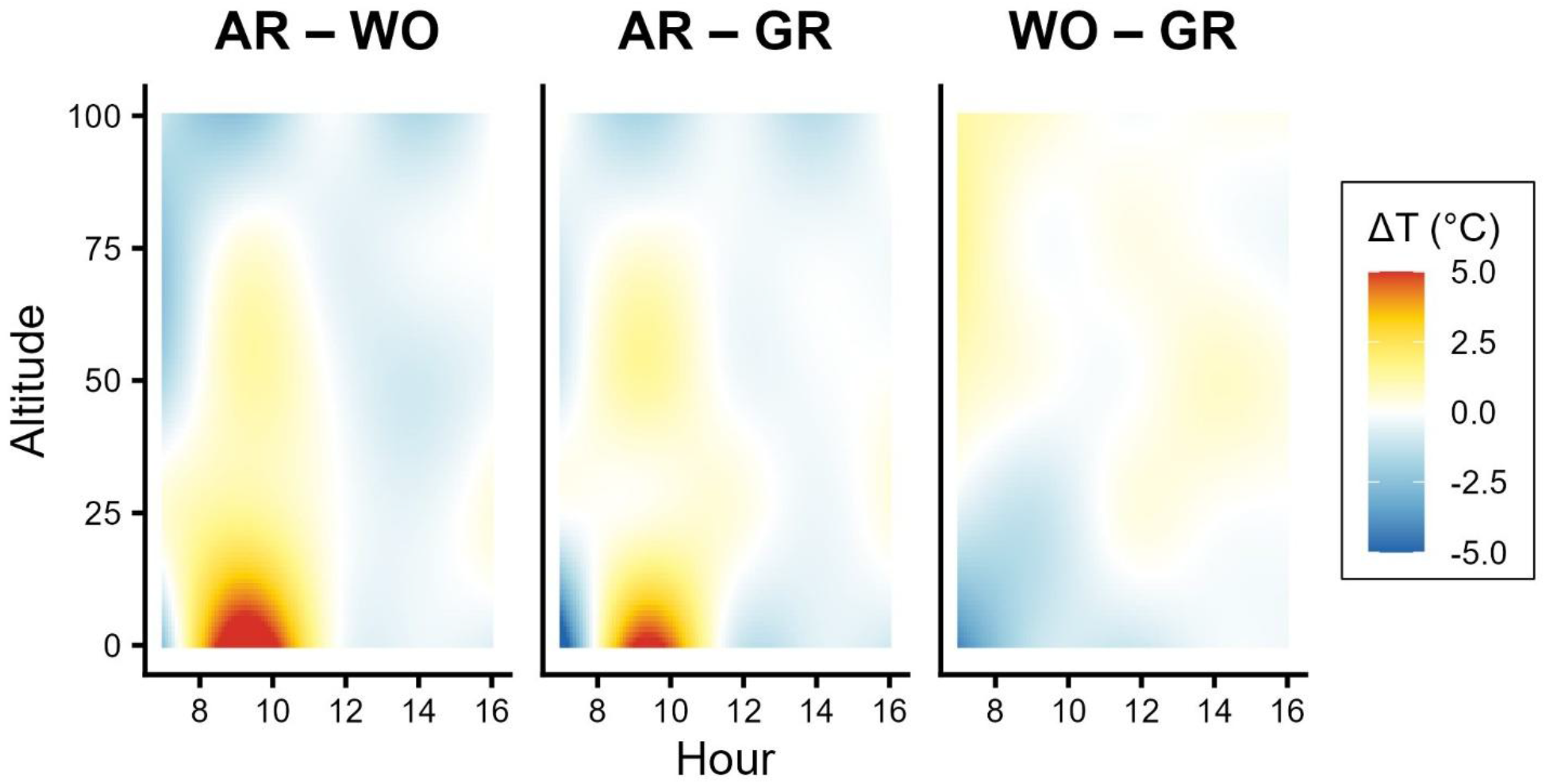
Pairwise differences in predicted air temperature (ΔT, °C) between land-cover classes as a function of hour of day and altitude, derived from the land-cover-specific smooths of the GAMM. Positive values indicate higher air temperature in the first land-cover class of each comparison (i.e. AR warmer than WO in AR – WO, AR warmer than GR in AR – GR, and WO warmer than GR in WO – GR), whereas negative values indicate higher temperature in the second class. The strongest differences were observed between artificial and vegetated surfaces, particularly during morning hours and at low altitudes. Differences between wooded vegetation and grass-covered areas were smaller but suggested slightly stronger cooling over wooded areas at low altitude

The non-linear effects of distance from vegetated areas were statistically significant for both wooded vegetation and grass-covered areas (s(Distance from trees): edf = 6.83, p < 0.001; s(Distance from grass): edf = 4.26, p < 0.001). In both cases, increasing distance from vegetated areas was associated with higher air temperatures at lower altitudes (Fig. 6).

**Fig. 6.**
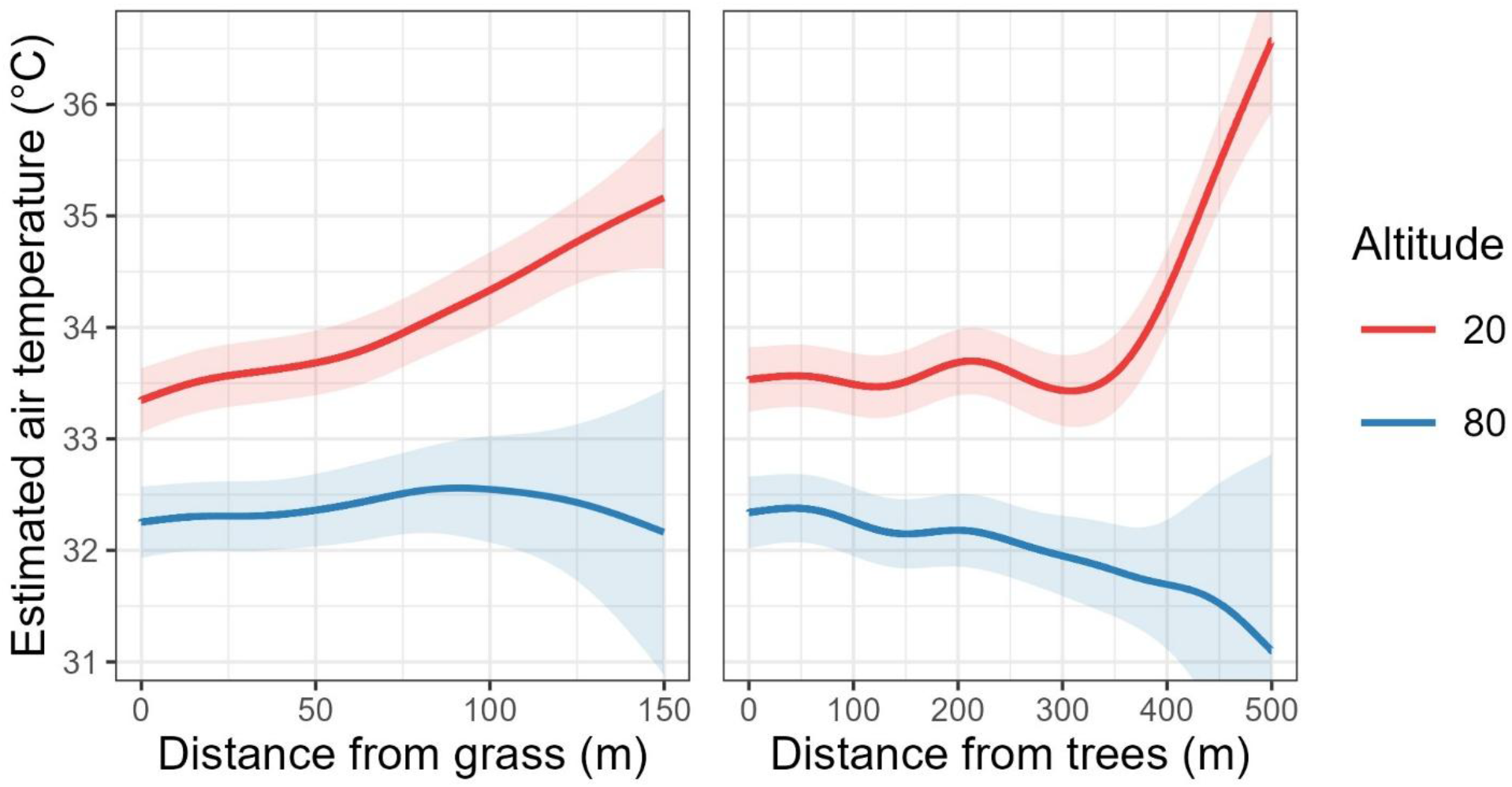
Interaction between altitude and distance from green areas (grass and trees) on predicted air temperature derived from the final GAMM. Predicted air temperature is shown as a function of distance from grass-covered areas (left panel) and wooded areas (right panel) at two representative altitudes (20 m and 80 m). Solid lines represent model predictions and shaded areas indicate 95% confidence intervals

The three-way interaction between time of day, altitude, and land-cover class was also significant (te(Hour, Altitude) by Land Cover: edf = 18.80–23.37, p < 0.001) (Fig. 7). This interaction revealed a general diurnal temperature pattern, characterized by a gradual increase until midday followed by a decrease during the afternoon. The magnitude and timing of this pattern varied across land-cover classes and elevation levels. At lower altitudes (20 m in Fig. 7), artificial surfaces were associated with higher air temperatures, particularly during the morning hours, whereas wooded and grass-covered areas maintained lower temperatures during the same period. These differences progressively diminished from midday to late afternoon and at higher altitudes (80 m in Fig. 7), suggesting that land cover and the diurnal cycle jointly modulate the intensity of urban thermal contrasts.

**Fig. 7.**
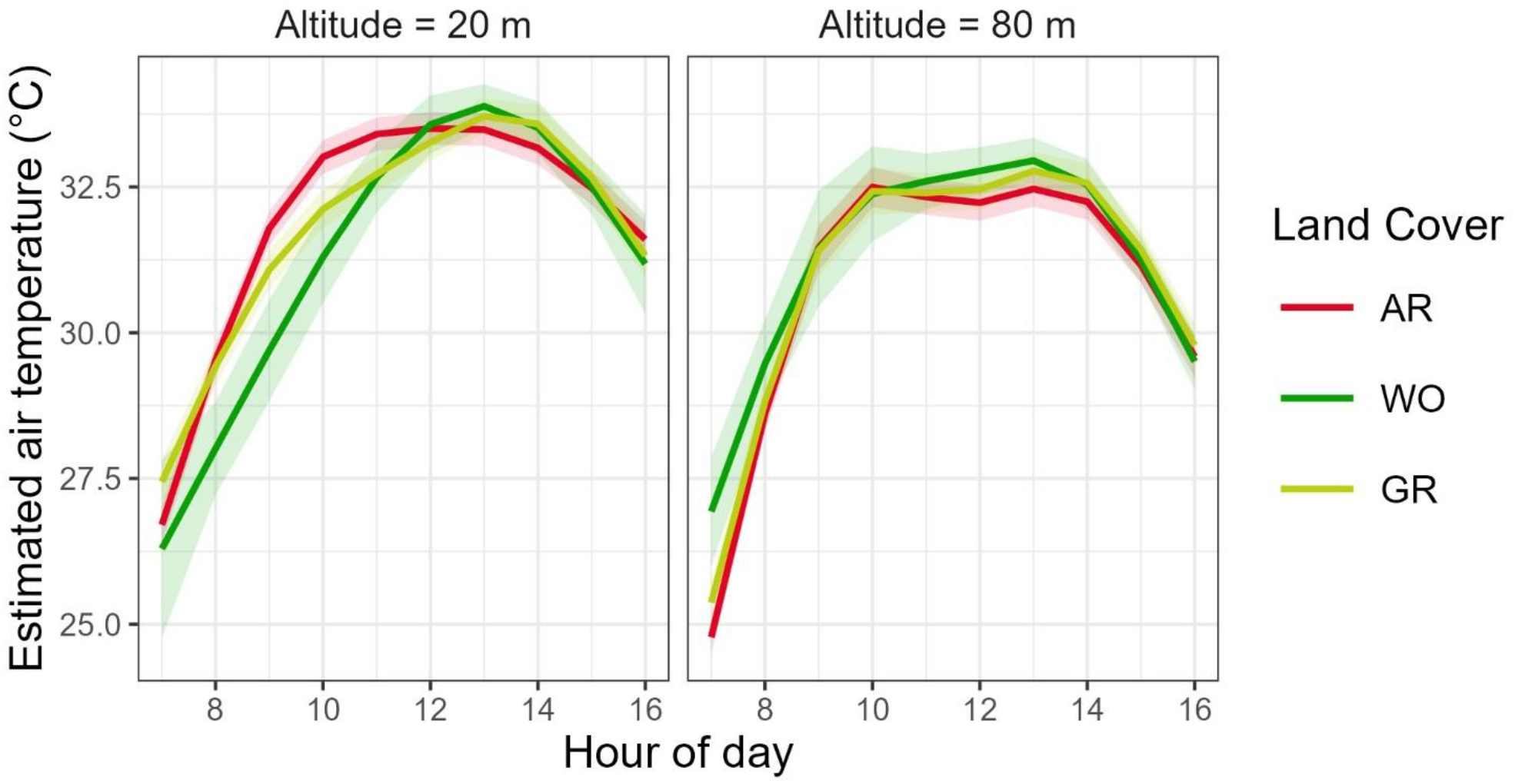
Predicted diurnal variation of air temperature across land cover classes at two representative altitudes. Modelled air temperature is shown as a function of hour of day for artificial surfaces (AR), wooded vegetation (WO), and grass vegetation (GR) at 20 m (left panel) and 80 m (right panel) elevation. Solid lines represent GAMM predictions and shaded areas indicate 95% confidence intervals

The random effect associated with sampling date was statistically significant (s(Date): edf = 10.0, F = 4153.4, p < 0.001), indicating substantial residual temporal variability in air temperature. This supports the inclusion of date as a random effect to account for day-to-day atmospheric conditions not explicitly captured by the fixed environmental covariates.

### 3.2 Heat mapping scenarios

Predictions from the GAMM were applied to a 100 x 100 m grid covering the study area to describe model-predicted air temperature over the city at 20 m above the ground during an average summer day at three representative times: 9:00, 12:00, and 15:00 (Fig. 8).

**Fig. 8.**
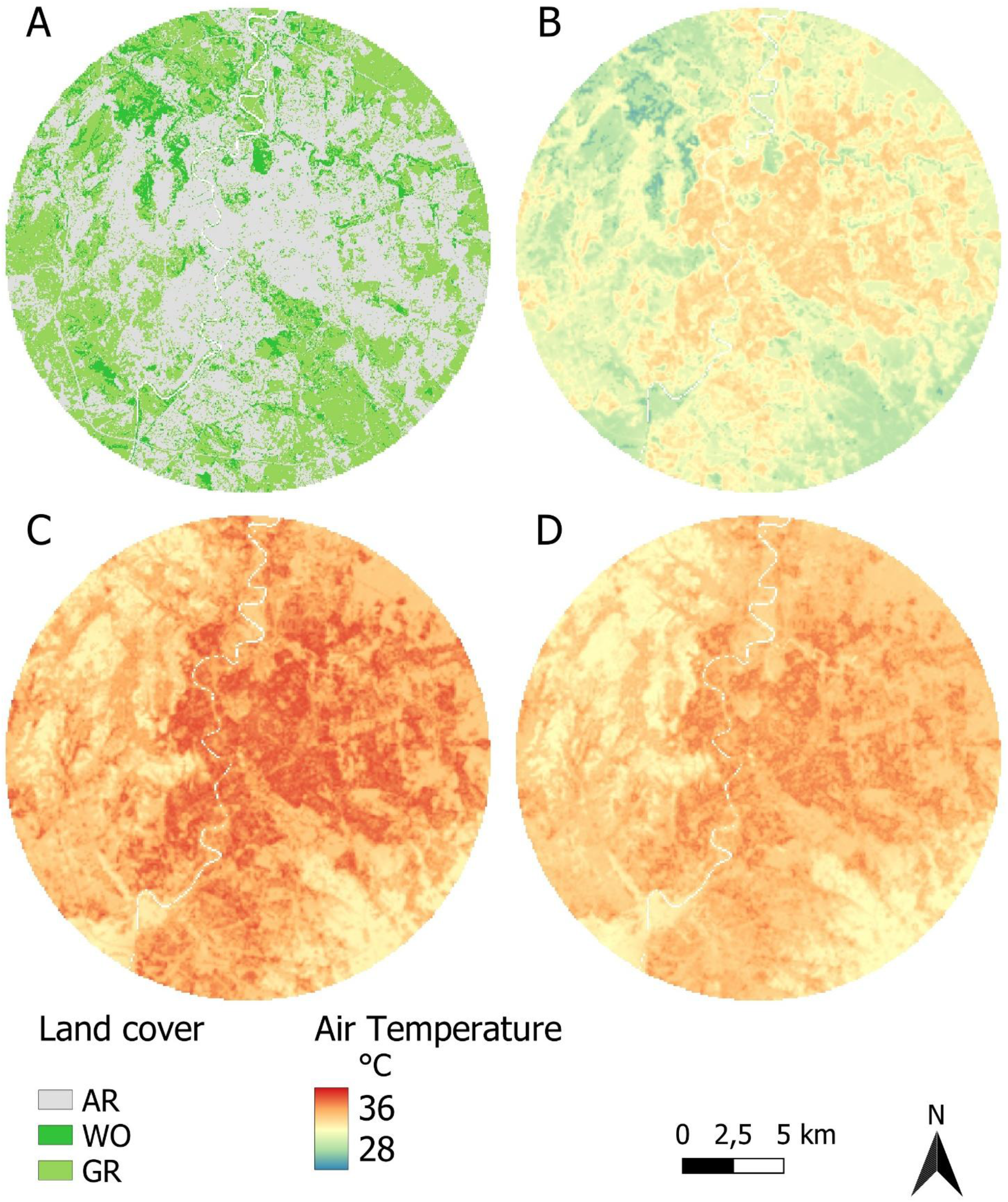
Spatial predictions of summer air temperature over Rome derived from the final GAMM. Air temperature was predicted at 20 m elevation on a 100 × 100 m grid. (a) Land cover classification showing artificial surfaces (AR), wooded vegetation (WO), and grass vegetation (GR). (b–d) Predicted air temperature (°C) at 09:00, 12:00, and 15:00

The mean (± SD) near-surface air temperature over the city was 32.1 ± 1.8 °C at 09:00, 35.3 + 1.5 °C at 12:00, and 34.4 + 1.4 °C at 15:00. Air temperatures displayed marked spatial heterogeneity across the urban domain at all analysed hours. In all scenarios, higher air temperatures were generally associated with areas characterised by artificial land cover, except at 12:00 when AR and WO displayed same average temperature (Table 2).

**Table 2.** Summary statistics of predicted air temperature (°C) at 20 m elevation for three representative summer hours (09:00, 12:00, and 15:00) across land cover classes (AR, WO, GR)

| Hour | Land cover |  |  |  |  |  |  |  |  |  |  |  |
| --- | --- | --- | --- | --- | --- | --- | --- | --- | --- | --- | --- | --- |
|  | AR |  |  |  | WO |  |  |  | GR |  |  |  |
|  | mean | max | min | sd | mean | max | min | sd | mean | max | min | sd |
| 09:00 | 33.5 | 34.4 | 32.1 | 1.4 | 30.7 | 31.4 | 29.8 | 1.3 | 32.2 | 32.9 | 31.2 | 1.4 |
| 12:00 | 35.3 | 36.2 | 33.9 | 1.4 | 35.3 | 36.0 | 34.4 | 1.3 | 34.3 | 35.1 | 33.4 | 1.4 |
| 15:00 | 34.3 | 35.2 | 32.9 | 1.4 | 33.6 | 34.4 | 32.7 | 1.3 | 33.8 | 34.6 | 32.8 | 1.4 |

Predicted temperatures increased from 9:00 to 12:00 and decreased by 15:00, with the highest values recorded at 12:00 across land-cover classes. At 9:00, the largest difference was between AR and WO, with AR being 2.8 °C warmer. At 12:00, differences among land-cover classes were smaller, reaching approximately 1 °C, and increased again by 15:00 (Fig. 8).

The spatial overlap between thermal hotspots, defined as cells above the 90^th^ percentile of predicted temperature, and areas with scarce vegetation, defined as cells with less than 10% local green cover, highlighted priority zones for mitigation planning.

HT_LGREEN areas had morning temperatures above 34.1 and covered a total of 21.9 km^2^, accounting for 4.8% of the total study area (Fig. 9).

**Fig. 9.**
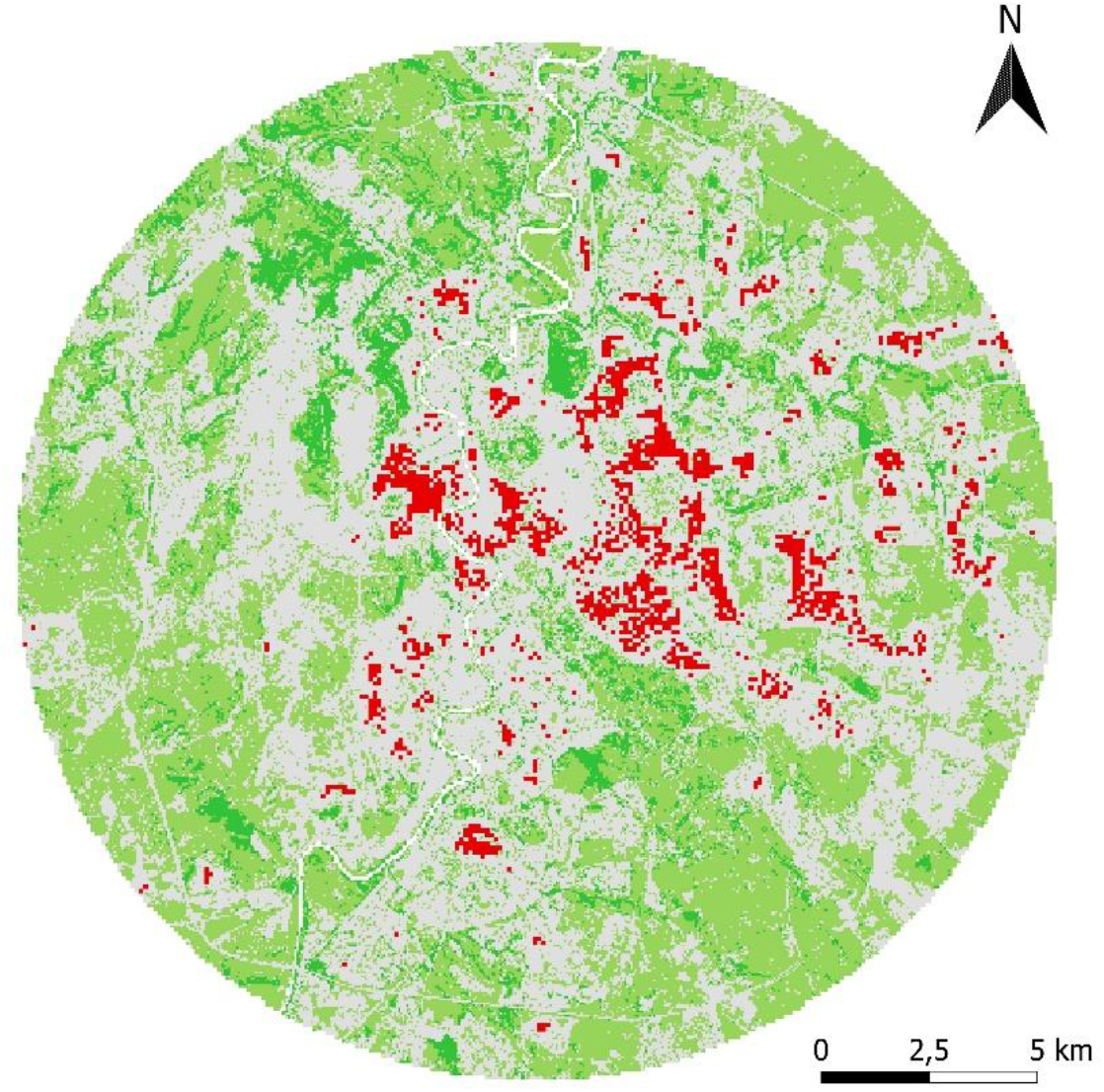
Priority areas for greening interventions within the study area. Areas with higher temperature and reduced green coverage are shown in red

## 4. Discussion

In this study, we demonstrated the potential of pigeons equipped with GPS–temperature sensors as mobile sentinels for monitoring fine-scale urban microclimates. We quantified summer thermal variability across the lower urban atmosphere and showed that air temperature varied markedly during the day, across land-cover classes, and at different altitudes.

These results allowed to identify hot urban zones in which targeted implementation of green infrastructure could help mitigate elevated air temperatures, with potential benefits for thermal comfort and long-term climate adaptation.

### 4.1 Use of pigeons

In our dataset, more observations were recorded over artificial and grass-covered areas, while fewer data points were collected over wooded areas. This pattern partially reflects the limited extent of wooded surfaces within the study area but may also be influenced by pigeon navigational behaviour. Previous GPS-based studies have shown that homing pigeons rely strongly on visually salient landscape features, such as edges and linear landmarks, during navigation (Lipp et al. 2004; Mann et al. 2011). Consequently, flight paths may be less frequent over large, visually homogeneous areas, including continuous forest patches, than over more heterogeneous or edge-rich urban environments.

In addition to navigational behaviour, the lower frequency of observations over wooded areas may also reflectpredator-avoidance behaviour. Urban and peri-urban forests can host raptors (Mirski 2020), which represent a significant predation risk for pigeons.

Despite these constraints in this study the use of pigeons as mobile sensing platforms proved useful for exploring the thermal complexity of urban environments. This approach enables direct sampling of air temperature along the lower atmospheric column, addressing some limitations of traditional observation systems.

To our knowledge, this study is the first to use bird-borne sensors to quantify daytime air-temperature variability across the urban atmosphere. Previous animal-based monitoring studies have primarily focused on nocturnal conditions or different taxa (Chudnovsky et al. 2023). In contrast, our approach provides direct measurements during daytime hours, which are particularly relevant for urban climate research because daytime temperatures are closely linked to human thermal exposure, outdoor activity patterns, and peak heat-stress conditions.

### 4.2 Urban green cooling

We found a clear association between urban vegetation and lower air temperature. Lower air temperatures were associated with increasing grass cover surfaces within the surrounding 500 m buffer, with predicted air temperatures decreasing progressively as the proportion of grass cover increased. Previous studies have shown that grass-covered surfaces can significantly reduce near-surface air temperature relative to artificial environments, particularly under clear and warm conditions (Bowler et al. 2010; Shashua-Bar et al. 2011). Our results suggest that continuous ground-level vegetation can moderate local thermal conditions even where tree canopy is sparse or absent. A similar same pattern was observed for wooded patches, with larger tree-covered areas associated with lower predicted air temperature. These cooling effects likely reflect the combined influence of evapotranspiration, higher surface albedo, shading, and reduced heat storage compared with artificial surfaces (Yilmaz et al. 2008).

Air temperature recorded along pigeon flight paths revealed pronounced spatial and temporal heterogeneity within the urban environment. Artificial surfaces consistently exhibited higher temperatures up to approximately 5 °C during the morning. This pattern is consistent with the key role of green spaces in mitigating urban heat (Gunawardena et al. 2017). Vegetated surfaces showed lower temperatures and greater variability, likely reflecting differences in energy exchange processes and heat storage capacity. Although differences between vegetated land-cover classes were generally modest, wooded areas tended to show slightly stronger cooling than grass-covered areas at low altitudes, below 25 m, especially during the early morning. This finding is consistent with the greater shading capacity, evapotranspiration and structural complexity of tree canopy (Armson et al. 2012). Overall, these patterns highlight how the urban mosaic generates distinct microclimates shaped by land-cover composition (Huang et al. 2008). The ability to detect thermal differences along flight paths also suggests that vegetation-related cooling effects extend beyond ground level into the lower urban atmosphere.

The cooling effect of green areas extended not only vertically but also horizontally, particularly at low altitudes. Air temperatures increased progressively with distance from both grass-covered and tree-covered areas, indicating a spatial cooling gradient extending beyond the green patches themselves. On average, air temperature increased by more than 3 °C at locations situated more than 300 m from wooded areas. For grass-covered areas the cooling gradient appeared more localised, with the strongest effect occurring within approximately 50 m. Although the cooling magnitude associated with herbaceous cover is typically lower than that provided by tree canopy, grass-covered areas can still play an important role in moderating neighbourhood-scale thermal conditions and improving outdoor thermal comfort. Overall, this pattern is consistent with previous studies showing that urban tree and grass cover reduces air temperature both locally and in surrounding areas (Aram et al. 2019). Our results are in line with Gallay et al. (2023), who reported buffer cooling effects associated with large grass-covered and tree-covered areas, with tree canopy exerting a stronger and more spatially extensive effect.

The three-way interaction between time of day, altitude, and land cover revealed a diurnal thermal cycle that varied with altitude. The cooling effect of vegetated areas was already evident during the morning hours and was particularly marked at low altitudes. At altitudes below 25 m, artificial surfaces were generally warmer, whereas tree-covered areas showed a wider thermal range and lower temperatures during the early part of the day. This pattern is consistent with the lower heat storage capacity and faster heat release of vegetated surfaces compared with artificial structures, which can contribute to reducing urban heat (Feyisa et al. 2014; Cai et al. 2023). The mitigating effect of tree vegetation relative to other land-cover types was evident during the morning hours and decreased around midday. Similar patterns have been reported in previous studies showing that the cooling effect of green areas can be particularly pronounced during hot summer conditions (Oliveira et al. 2011; Li and Norford 2016), which is especially relevant in Mediterranean cities. This temporal pattern is important for understanding the persistence of the urban heat island (Gartland 2012), as it suggests that temperature differences between land-cover classes are established before the daily maximum and tend to diminish when the broader urban area reaches high temperatures. Green spaces therefore play a critical role in maintaining cooler air within the near-surface urban layer, where human exposure occurs. By moderating temperatures in this zone during summer, urban vegetation contributes to reducing heat stress and improving thermal comfort and public health outcomes (Vicedo-Cabrera et al. 2021). Importantly, our results suggest that the cooling influence of vegetation is not confined to ground level, but propagates vertically throughout the lower urban atmosphere, reaching heights of up to approximately 75 m. This suggests that the benefits of urban greening are not limited to street-level conditions but may also influence the thermal environment of mid-and high-rise buildings. Such vertical cooling likely arises from a combination of reduced sensible heat flux, enhanced cooling by evaporation and vertical turbulent mixing that redistributes cooler air masses above vegetated surfaces.

From an applied perspective, this finding has relevant implications for urban design in densely built Mediterranean cities, where a large proportion of the population resides in multi-storey buildings. By extending cooling effects into the urban canopy layer, vegetation may contribute to lowering indoor temperatures, reducing energy demand for cooling, and improving thermal comfort not only outdoors but also within upper floors of buildings. This highlights the importance of incorporating vegetation not only as ground-level green space, but also as a three-dimensional component of urban climate mitigation strategies, including green corridors, urban forests, green roofs (Wang et al. 2022) and vertical greening systems (Susca et al. 2023).

### 4.3 Heat mapping and priority areas

Spatial predictions derived from the GAMM provided a fine-scale, model-based representation of air temperature variability within the urban canopy layer (Oke 1988), enabling the identification of localized thermal hotspots and potential priority areas for mitigation. Predicted temperatures showed consistent spatial heterogeneity across the study area, with the highest values associated with artificial surfaces and the lowest values occurring in areas characterized by tree-and grass-covered vegetation. This pattern was evident at all analysed hours, confirming the dominant influence of land cover on near-surface air temperature.

The cooling influence of vegetation is particularly relevant in Mediterranean cities, where intense solar radiation and prolonged dry periods can amplify daytime heating. By reducing early heat build-up, green spaces can help delay peak temperatures and lower cumulative heat exposure (Oliveira et al. 2011). For this reason, thermal hotspots were identified using morning air temperatures, allowing the detection of zones where early heat accumulation may increase the likelihood of prolonged thermal stress. Targeting mitigation interventions in these areas may be more effective than focusing on locations where peak temperatures occur later in the day. Such interventions could help delay daytime heat build-up and enhance nocturnal cooling, processes that play a critical role in the development and persistence of the urban heat island effect (Edmondson et al. 2016). The spatial identification of priority areas provides practical guidance for urban greening strategies aimed at improving thermal resilience (Turner-Skoff and Cavender 2019).

In our study, priority areas for mitigation represented approximately 4.8% of the study area and were spatially concentrated rather than uniformly distributed across the urban landscape. This suggests that targeted greening interventions, such as strategic tree planting (Gill et al. 2007), expansion of green corridors (Norton et al. 2015), and green roofs (Oberndorfer et al. 2007; Li et al. 2014), could provide local cooling benefits without requiring large-scale land-use changes. Increasing tree canopy and ground vegetation in these priority areas could reduce local air temperatures, enhance thermal comfort, and contribute to climate resilience. In Mediterranean cities, where extreme heat events pose significant risks to human health (Maggiotto et al. 2021) and are projected to increase (Zittis et al. 2016), spatially targeted nature-based solutions represent an effective strategy to mitigate urban heat island intensity and improve urban liveability. Recent studies also indicate that urban temperatures have risen steadily over the last 25 years and that the expansion of green areas can contribute to mitigating these trends (Mohammad et al. 2022).

In many densely built cities, however, the creation of new ground-level green areas, such as parks, or the implementation of strategic tree planting is often constrained by limited space availability (Haaland and van Den Bosch 2015), competing land uses, and the high economic costs associated with urban redevelopment. In this context, alternative nature-based solutions that can be implemented within the existing built environment become particularly important. Green roofs represent one of the most promising strategies for increasing urban vegetation cover without requiring additional land consumption (Susca et al. 2011). Therefore, integrating green roofs into urban planning policies may provide an effective and scalable approach to enhance thermal resilience in cities where horizontal expansion of green spaces is limited.

### 4.4 Possible limitations and future directions

Some limitations should be acknowledged. Data collection was restricted to daytime hours during the summer period, preventing the assessment of nocturnal thermal dynamics, which play a key role in nighttime cooling processes and in the persistence of the urban heat island. In addition, the analysis was conducted within a single metropolitan area, which may limit the transferability of the findings to cities with different climatic conditions and urban morphologies.

The use of homing pigeons as mobile sampling platforms also resulted in uneven spatial coverage, as flight trajectories were shaped by homing behaviour rather than by a systematic sampling design. This limits the interpretation of the predicted maps as absolute spatial temperature fields. However, this limitation is inherent to the exploratory nature of this study and could be addressed in future applications by increasing the number of tracked individuals and implementing releases from a more spatially distributed set of locations, thereby ensuring a more uniform sampling of the urban area.

Despite these constraints, sampling along the urban air column enabled the characterization of fine-scale three-dimensional temperature variability, addressing some limitations of traditional fixed stations and satellite-based measurements. Furthermore, the bio-assisted monitoring approach is relatively low-cost and logistically simple compared to conventional fixed meteorological networks, which require substantial installation and maintenance efforts. The cost of animal-borne sensors is also likely to decrease with ongoing technological development, enhancing the scalability and replicability of this approach for urban climate research.

An additional implication of this study concerns the societal perception of urban pigeons, which are often regarded as nuisance or pest species in many cities (Giunchi et al. 2012; Hermann et al. 2024). Our findings show that these animals can act as mobile environmental sentinels, collecting high-resolution climatic data in complex urban environments. This perspective highlights the potential to reframe an underappreciated urban species as a useful platform for environmental monitoring and climate adaptation research. Future studies could explore whether this approach can be extended to feral pigeon populations equipped with lightweight sensors, potentially enabling more continuous and spatially extensive data collection. Such applications would further enhance the scalability of bio-assisted monitoring systems and reinforce the integration of animal-based sensing into urban climate research and smart-city infrastructures.

A further methodological limitation relates to the maximum flight altitude typically reached by pigeons during routine movements within urban environments. In our study, most observations were concentrated in the lower portion of the urban canopy layer, generally below approximately 70 m. Although this altitude range is highly relevant for assessing human thermal exposure and street-level microclimates, it does not fully capture temperature dynamics at higher elevations within dense urban settings. Future applications could address this limitation by integrating bird species with different flight ecologies. For example, urban gulls are known to exploit vertical air currents and reach greater flight altitudes than pigeons (Corman and Garthe 2014). Equipping gulls or other high-flying urban birds with lightweight environmental sensors could therefore extend the vertical coverage of bio-assisted monitoring systems, enabling a more comprehensive characterization of temperature gradients, or other environmental variables, throughout the entire urban boundary layer. Furthermore, the nocturnal activity of some gull species (Fernandes et al. 2026) could provide information on nighttime thermal dynamics, complementing the daytime measurements obtained from pigeons.

## 5. Conclusions

Our findings highlight the potential of animal-borne sensors to characterise three-dimensional urban microclimates, while also showing the important role of urban vegetation in buffering summer air temperatures within the lower urban atmosphere. More broadly, this study supports the spatial prioritisation of nature-based solutions in large metropolitan areas. Although the study focused on Rome, the patterns identified, including early-morning warming of built-up areas, cooling gradients associated with proximity to trees, and the vertical attenuation of land-cover effects, are likely relevant to other Mediterranean and temperate cities with comparable urban morphology (Schwaab et al. 2021).

Overall, effective urban heat mitigation requires spatially targeted strategies that consider land-cover composition together with the temporal dynamics of urban temperatures. Identifying areas prone to early heat accumulation can provide useful information for climate adaptation planning and support the prioritisation of greening interventions to enhance urban thermal resilience. This study therefore provides evidence for climate-resilient urban design and highlights the potential of dynamic bio-assisted sensing for monitoring urban environments.

## Data availability

The data supporting the findings of this study are available from the corresponding author upon reasonable request.

## Author contributions

Giacomo Grosso: Conceptualization, Data curation, Formal analysis, Investigation, Methodology, Writing – original draft. Luca Nelli: Formal analysis, Writing – review and editing. Claudio Carere: Supervision. Giacomo Dell’Omo: Resources, Supervision, Writing – review and editing. All authors read and approved the final manuscript.

## Use of generative AI in manuscript preparation

During the preparation of this work the authors used ChatGPT (OpenAI) and Claude (Anthropic) to improve language quality, structure sentences, and assist in drafting some parts of the manuscript.

## Acknowledgments

We gratefully acknowledge the support of the *Ornis italica* Association for logistical assistance and access to the homing pigeon facilities used during the study. We thank Matteo Trivellini and Valeria Jennings for technical support and assistance during field activities.

