## Supplementary file for "Bird-assisted sampling of air temperature reveals three-dimensional cooling effects of urban vegetation"

*Corresponding authors:*

*Giacomo Grosso*

*ORCID ID: 0009-0004-8543-5288*

| Date | N of valid GPS points |
|---|---|
| 01/07/2023 | 298 |
| 01/07/2024 | 1371 |
| 03/07/2021 | 4554 |
| 04/07/2024 | 1399 |
| 15/06/2024 | 3753 |
| 16/06/2023 | 4377 |
| 17/06/2025 | 2128 |
| 18/06/2025 | 726 |
| 23/06/2024 | 1873 |
| 24/06/2023 | 3624 |
| 26/06/2024 | 2177 |

**Table S1.** Number of valid GPS records retained for analysis for each sampling date after applying all data filtering procedures.

*Parametric coefficients*

| Term | Estimate | Std. Error | t value | p-value |
|---|---|---|---|---|
| Intercept | 31.680 | 0.780 | 40.83 | <0.001 |
| DEM | −0.01220 | 0.000 | −32.48 | <0.001 |
| % Semi-natural non-forest (500 m) | −0.01143 | 0.001 | −20.32 | <0.001 |
| Flight speed | −0.01260 | 0.001 | −15.52 | <0.001 |
| Tree-covered area (ha) | −0.00142 | 0.001 | −2.71 | 0.007 |

*Smooth and interaction terms*

| Term | edf | Ref.df | F | p-value |
|---|---|---|---|---|
| s(Distance from trees) | 6.83 | 9 | 6.03 | <0.001 |
| s(Distance from grass) | 4.26 | 9 | 2.41 | <0.001 |
| ti(Altitude, Distance from trees) | 15.29 | 16 | 66.86 | <0.001 |
| ti(Altitude, Distance from grass) | 11.40 | 16 | 9.04 | <0.001 |
| te(Hour, Altitude): Land cover (ART) | 23.37 | 24 | 2611.11 | <0.001 |
| te(Hour, Altitude): Land cover (WO) | 18.80 | 24 | 117.42 | <0.001 |
| te(Hour, Altitude): Land cover (GR) | 22.66 | 24 | 846.15 | <0.001 |
| s(Date) | 10.00 | 10 | 4153.40 | <0.001 |

*Tab. S2. Parametric, smooth and interaction terms of the best model. Summary of parametric coefficients and approximate significance of smooth and interaction terms of the generalized additive model (GAM) fitted to air temperature data. For smooth terms, effective degrees of freedom (edf), F statistics and p-values are reported. Ref.df (reference degrees of freedom) represents the degrees of freedom used to compute the approximate F-statistics for smooth and interaction terms in the GAM.*

Model performance

| Metric | Value |
|---|---|
| Adjusted $R^2$ | 0.73 |
| Deviance explained (%) | 73.50 |
| Scale estimate | 2.26 |
| Estimation method | fREML |
| Sample size (n) | 26280 |

*Tab. S3. Model performance. Overall performance metrics of the generalized additive model, including adjusted $R^2$, percentage of deviance explained, scale estimate (corresponding to the residual variance of the model), estimation method and umber of observations.*

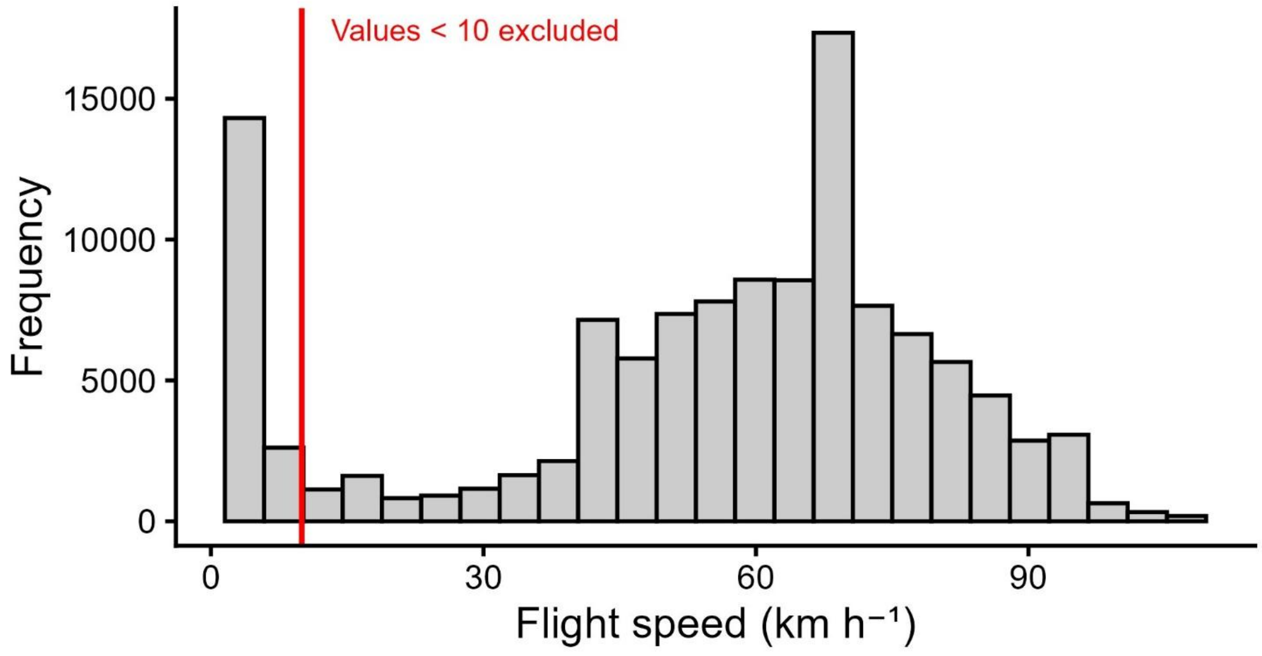


Fig. S1. Distribution of recorded flight speeds (km $h^{-1}$) from GPS-tracked pigeons before filtering. The red vertical line indicates the threshold used to exclude low-speed records (<10 km $h^{-1}$), which were removed from the analysis.

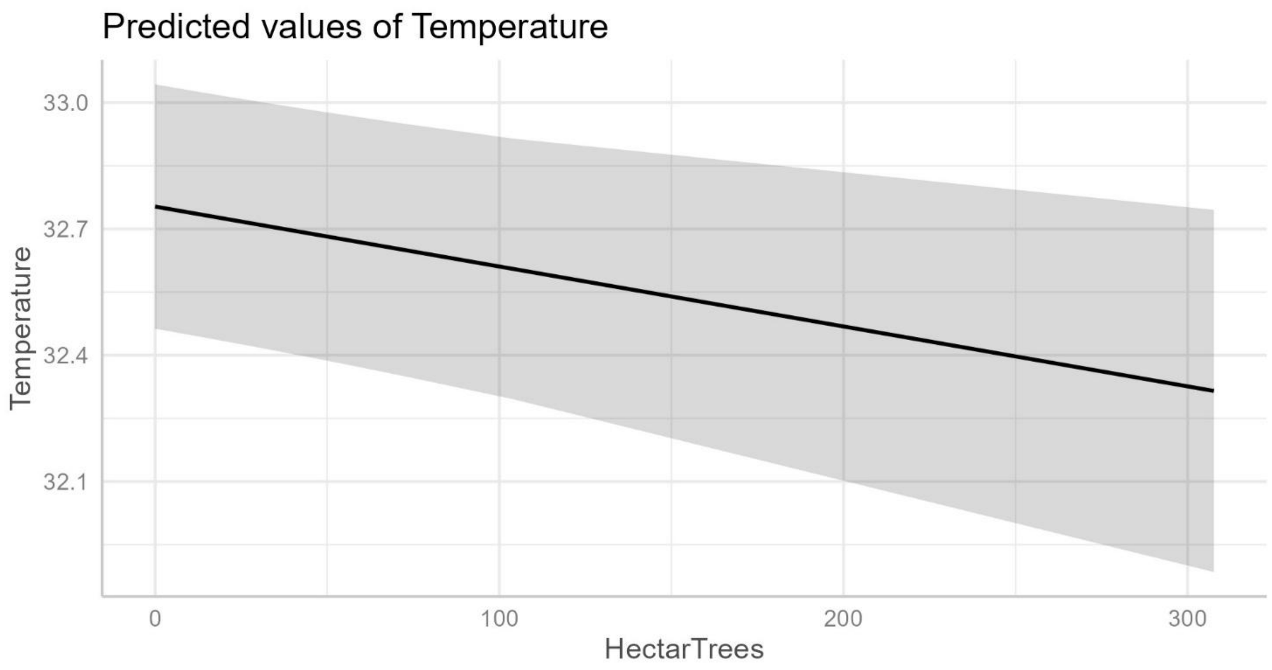


Figure S2. Partial effect of tree-covered area (ha) on predicted air temperature derived from the optimal GAMM. Predicted air temperature decreases with increasing extent of tree-covered area, indicating a cooling effect associated with larger wooded patches.

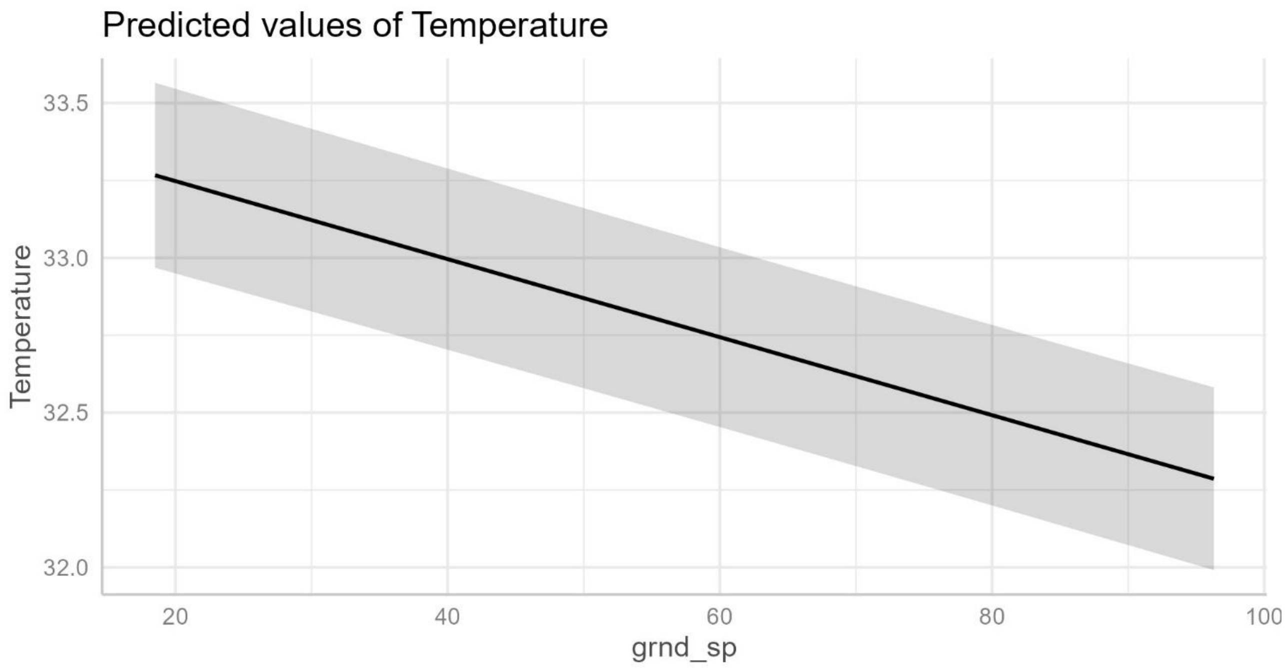


Figure S3. *Partial effect of* ***flight speed (km h⁻¹)*** *on predicted air temperature derived from the optimal GAMM. Predicted air temperature decreases with increasing flight speed.*

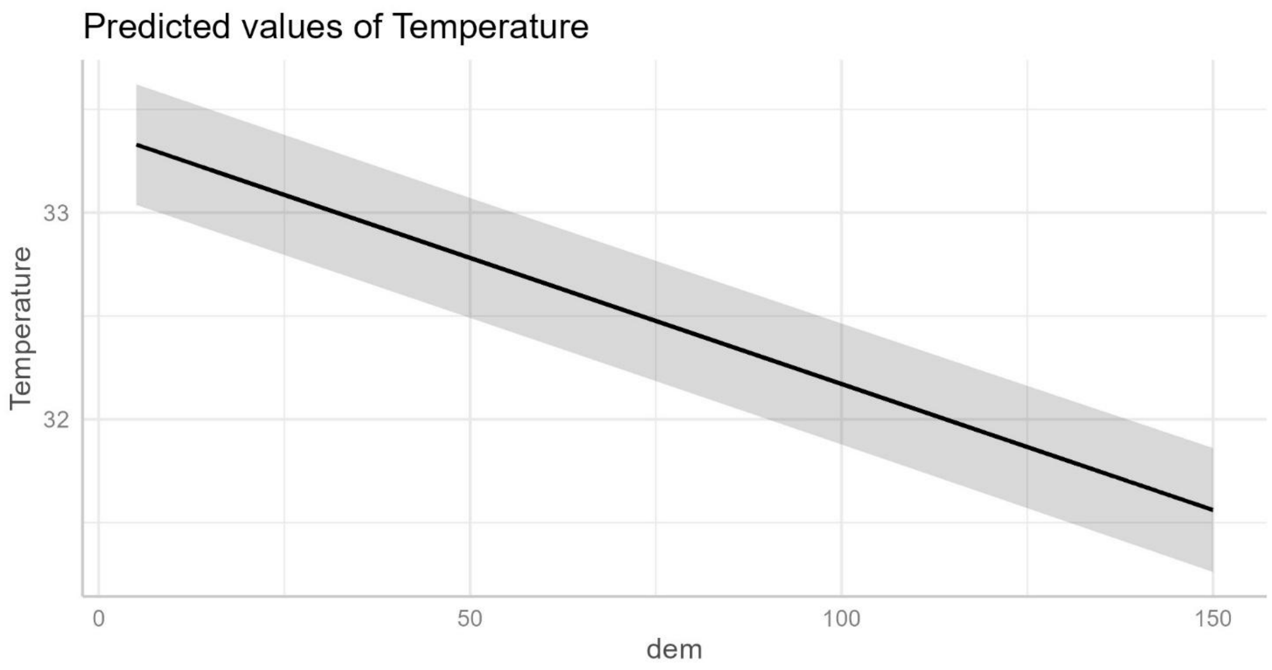


*Figure S4.* Partial effect of **digital elevation model (DEM)** on predicted air temperature derived from the optimal GAMM. Predicted air temperature decreases with increasing elevation.
